# Diversity in Clinical and Biomedical Research: A promise yet to be fulfilled

**DOI:** 10.1101/034538

**Authors:** Sam S. Oh, Joshua Galanter, Neeta Thakur, Maria Pino-Yanes, Nicolas E. Barcelo, Marquitta J. White, Danille M. de Bruin, Ruth M. Greenblatt, Kirsten Bibbins-Domingo, Alan H.B. Wu, Luisa N. Borrell, Chris Gunter, Neil R. Powe, Esteban G. Burchard

**Author notes:** These authors contributed equally to this manuscript (Neil R. Powe, Esteban G. Burchard).

## Abstract

**Summary Points:**

- Health disparities persist across race/ethnicity for the majority of *Healthy People 2010* health indicators.
- Most physicians and scientists are informed by research extrapolated from a largely homogenous population, usually White and male.
- A growing proportion of Americans are not fully benefiting from clinical and biomedical advances since racial and ethnic minorities make up nearly 40% of the U.S. population.
- Ignoring the racial/ethnic diversity of the U.S. population is a missed scientific opportunity to fully understand the factors that lead to disease or health.
- U.S. biomedical research and study populations must better reflect the country’s changing demographics. Adequate representation of diverse populations in scientific research is imperative as a matter of social justice, economics, and science.

---

In 1993, the National Institutes of Health (NIH) Revitalization Act was passed by United States (U.S.) Congress and signed into law by President Clinton [1]. The Act called for the NIH to require that all federally-funded clinical research prioritize the inclusion of women and minorities and that research participant characteristics be disclosed in research documentation [1]. When pivotal NIH-funded studies included large proportions of women by design, they made important, clinically relevant scientific contributions by identifying sex-specific differences in symptoms, pathologies, and treatment response [2-4]. In continuation of this effort, the NIH announced new measures to enhance gender equity [5]. Herein, we evaluate the impact of the Revitalization Act’s other stated aim: diversifying study populations by race/ethnicity. We also make suggestions on what we believe will bolster the Revitalization Act’s effect in shaping clinical and biomedical research and thereby provide guidance for President Obama’s new Precision Medicine Initiative (PMI) [6].

## Disease pattern, clinical presentation, and therapeutic response can vary dramatically by race/ethnicity and ancestral background

Race is a social construct rooted in cultural identity and shaped by historic and current events, which influence an individual’s behavior and place of residence. Genetic variation correlates with self-identified race [7], and this genetic variation also correlates with clinical presentation and therapeutic response. Thus, while not every study needs to examine racial differences or include all racial/ethnic groups, we feel that the group(s) included should be representative of their larger population(s) such that including an adequate proportion of racially/ethnically diverse groups in clinical and biomedical research can provide meaningful opportunities to examine the complex relationship of ancestral influences, environmental exposures, and social factors. In turn, understanding the interaction between the social and environmental milieu with an individual’s genomic profile and genetic ancestry can extend our understanding of disease pathology and expand therapeutic options for everyone [8]. For example, up to 75% of Pacific Islanders are unable to convert the antiplatelet drug clopidogrel into its active form and are at higher risk for adverse outcomes following angioplasty [9,10]. Other examples are listed in Table 1.

**Table 1.**
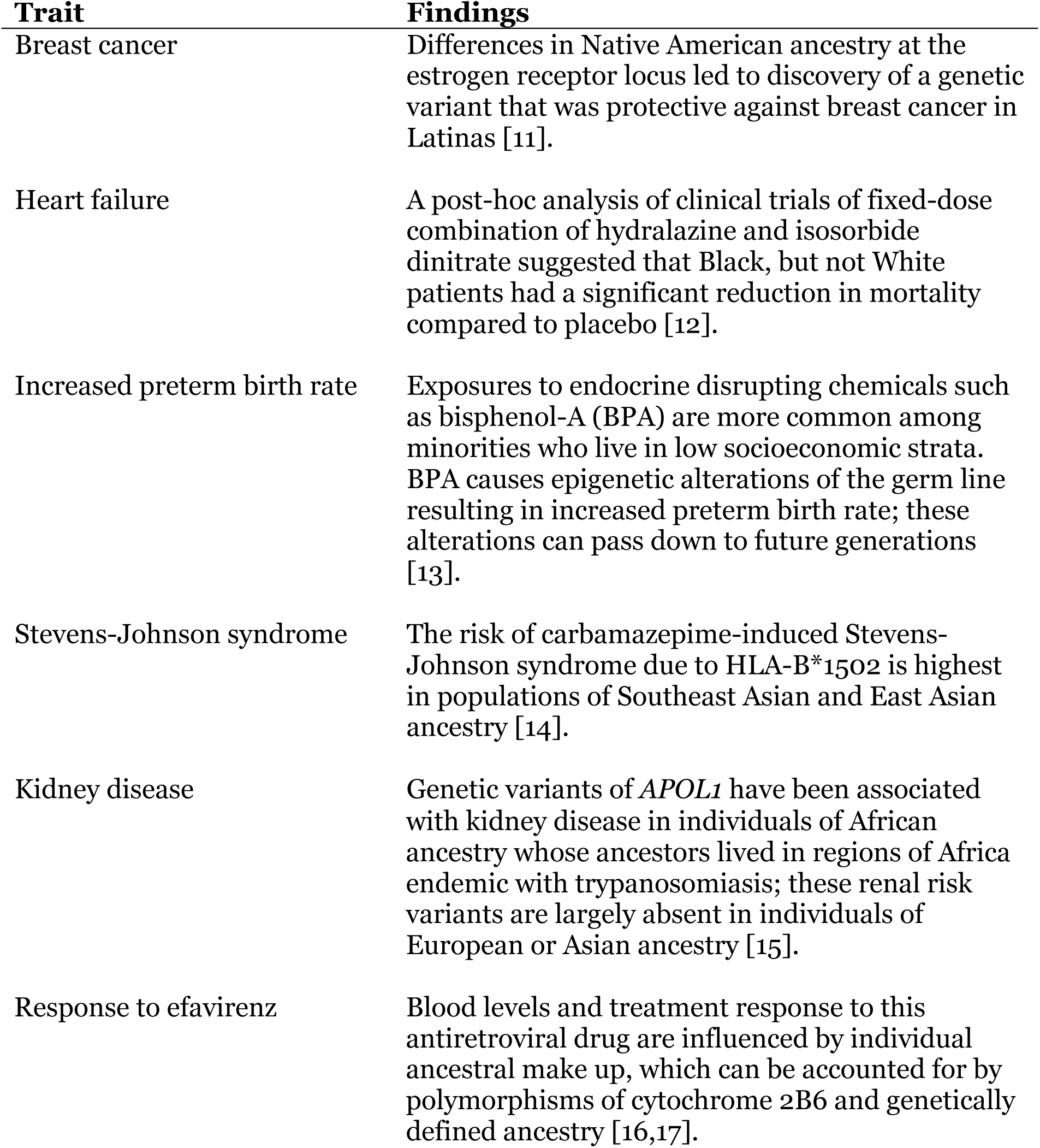
Insights from studies conducted in diverse race/ethnic groups.

## Past research has under-studied minorities

The U.S. has been regarded as a “global lead” and “exemplar” in biomedical and clinical health research since the end of the Cold War [18]. Yet, few U.S. biomedical studies focus recruitment efforts on attaining adequate minority representation nor their research attention to factors most relevant to minority health [19]. Since the passage of Revitalization Act in 1993, less than 2% of more than 10,000 cancer clinical trials funded by the National Cancer Institute included enough minority participants to meet the NIH’s own criteria and goals [20]. Moreover, less than 5% of NIH-funded respiratory research reported inclusion of racial/ethnic minorities [21]. Minority enrollment in cancer clinical trials remains inadequate despite striking racial/ethnic disparities in cancer incidence and mortality [22,23]. Similar incongruities between disease burden and representation in biomedical research exist for cardiovascular diseases and diabetes [24,25]. These disparities have economic consequences: eliminating racial/ethnic health disparities would have reduced total medical costs during 2003–2006 by more than $1.2 trillion [26]. Some NIH reviewers have argued that the inclusion of diverse groups will increase the financial costs of clinical and biomedical research. However, it is generally agreed upon that the long-term financial benefits outweigh short-term expenses [27]. The social, biomedical and economic costs of inaction are ameliorated by a new appreciation for the clinical and biomedical benefits achieved through precision medicine when applied to all populations [6]. The proportion of taxpayers who have not gained optimal benefit from scientific discoveries they are funding continues to grow with the changing U.S. demographics. Therefore, ensuring that diverse populations are adequately included in scientific research is imperative not only in terms of scientific integrity and fiduciary responsibility, but also as a matter of social justice.

## Barriers to diversify research need concerted attention

While U.S. minorities may be as willing to participate in health research as non-Hispanic whites [28], barriers to participation among minority populations must be addressed and will require buy-in from stakeholders: funders, academic institutions, investigators, and potential research participants [29]. Minority populations often have limited access to specialty care centers that serve as referral sources for clinical studies, resulting in a lack of an effective referral base [30]. Other barriers include, but are not limited to, fears of exploitation in medical research [31], financial constraints [32], competing demands of time, lack of access to information and comprehension about research, unique cultural and linguistic differences, fears of unintended outcomes, stigmatization, and health care discrimination [31].

Highly feasible changes can increase minority participation despite the challenges described. Ideally, investigators would reflect the communities being studied. Given the tremendous disparities in our biomedical workforce, we must seek out other realistic solutions. For example, some participants prefer studies that include research staff who share their same culture and with whom they can communicate in their own language [31]. Potential contributors are also more likely to partake when recruited by research staff they personally know or with whom they identify [33,34]. Town hall meetings and study newsletters can be adapted to the language and reading level requirements of target groups; these can describe how collected data will be used, ensuring transparency and allaying fears stemming from lack of information [29]. Challenges of transportation, childcare, work hour considerations, and meals can be addressed via payment, travel support, flexible recruitment hours and locations, provision of food during study visits, and positioning study sites in areas with diverse residents. To compensate for the limited internal referral base, tertiary care centers can partner with community healthcare providers. Targeted advertising (e.g., on public transportation) can reach potential participants at a moderate cost. Nonetheless, outreach and external partnerships introduce costs and effort that can raise recruitment budgets. The Revitalization Act specifically prohibits cost considerations from being a reason to exclude minorities, and NIH study sections are instructed to disregard budgetary requests in evaluating a project’s scientific merit. However, our experience in grant reviewing has been that in practice, the size of budgetary requests can bias reviewers. Grant applicants, in turn, react by submitting proposals with inadequate budgets to recruit minority participants so as not to “raise eyebrows” of reviewers.

Minorities would likely to be as willing to be involved in research as whites if problems of diversity could be better addressed. Some of these problems may stem from issues within the research community and its own profound diversity gap. Minority physicians and scientists are more likely to conduct research in minority populations and are often best suited to gain the trust of minority communities, but they are also significantly underrepresented in medical and scientific communities [35]. For example, Blacks or African Americans and Hispanics respectively represented only 4.3% and 7.2% of doctorate degree awardees in biomedical sciences in 2013 although they represented 13.2% and 17.1% of the U.S. population during the same period [36]. Moreover, less than 2% of NIH principal investigators on research project grants are Black [37], a proportion much lower than in the general U.S. population (10.2%) [38]. Similar disparities were observed for Latinos (3.4% vs. 12.5%), American Indians and Alaska Natives (0.4% vs. 0.7%), and Native Hawaiians and Other Pacific Islanders (1.2% vs. 10.2%).

To further complicate the picture, an NIH study of research grant awards found that the proportion of applications funded was 13% lower for Blacks or African Americans and 4% lower for Asians than among Whites [39]. According to demographic information provided by the NIH’s Office of Extramural Research under the Freedom of Information Act, the award rate for R01 or equivalent grants has been consistently lower among non-White applicants (Pacific Islander, Native Hawaiian, African American, American Indian, and Asian) than White applicants (42.1% versus 48.6% in 1985 and 19.3% versus 23.3% in 2013) [40].

Contributors to funding disparities arise throughout the research application review process [41]. The NIH has commented on reviewer bias [42], acknowledging that the probability of funding after peer review does not differ by race, but that minority investigators tend to receive lower priority scores from peer review, indicating that the review process is biased against applications from minority investigators. The relative absence of minority participants throughout the research application evaluation process may contribute to this problem, since underrepresented minorities comprised 10% of NIH study section reviewers in 2000 and only 10.9% in 2013 [40]. Increasing minority representation within the research community could in itself promote better science. Diverse research teams are more likely to have diverse ideas [43], which may explain why manuscripts authored by multi-ethnic research teams are more likely to be cited than publications authored by authors of the same ethnicity [44]. However, since study section members are drawn from the pool of successfully funded researchers, funding disparities have a self-perpetuating effect [45] and functionally eliminate scientists best suited to respond to the call to action we describe.

## How can the NIH (re)catalyze diversity in research?

The Revitalization Act intended to re-catalyze diversity in biomedical research by increasing minority representation. President Obama’s Precision Medicine Initiative plans to enroll a cohort of one million or more Americans that will provide the platform for expanding our knowledge and benefit the nation for many years to come. It is time to heed to the President’s call to action given the changing U.S. demographics. The NIH should be empowered to set and enforce recruitment of diverse research populations as the default and require scientific justification for limited or selected study population enrollment, as they have just set policies to do for sex balance [5]. Other U.S. government agencies (e.g., Centers for Disease Control and Prevention, Food and Drug Administration, Agency for Healthcare Research and Quality, Patient-Centered Outcomes Research Institute, Department of Defense) should be similarly empowered. Recruitment approaches should be formally included as criteria for scientific merit scoring, rather than the current application of such criteria after scoring.

In this vein, the NIH should include race/ethnicity as a criterion for assigning priority scores to ensure that well-characterized cohorts and clinical trials not only answer questions relevant to the growing diversity of the U.S. population but are also appropriately statistically powered. The same techniques for monitoring sex/gender inclusion [5] should be used to explicitly review minority accruals over the course of the award, and adjust funding levels accordingly. We believe this would prompt researchers of all racial/ethnic and cultural backgrounds to incorporate understudied populations in their research studies.

To their credit, the NIH is actively addressing many of the issues we have mentioned. Following President Obama’s PMI announcement during his 2015 State of the Union address, the NIH has actively solicited feedback [46] to help guide creation of a diverse research cohort of 1 million or more Americans [47]. The NIH has since hosted several workshops to develop a vision for building the national PMI cohort, and maximizing cohort diversity (across socioeconomic standing, geography, sexual orientation, education, and age, in addition to race/ethnicity) has been an ongoing topic at these workshops [48]. In particular, participant and public engagement, diversity and inclusion, and health disparities considerations for the development of a national research cohort were among the topics discussed at a workshop dedicated to participant engagement and health equity [49].

We applaud and encourage the NIH’s focus on diversifying the makeup of the forthcoming PMI cohort. To build on these efforts, an administrative supplement for currently funded research to investigate racial/ethnic differences in health and therapeutics should be created, similar to efforts by the Office of Research on Women’s Health to promote discovery of sex differences [50]. This supplement would be hypothesis-generating and show the NIH’s commitment to diversify study populations throughout all Institutes. The NIH should also incentivize collaboration amongst groups with similar approaches and data elements so that adequately powered analyses can examine racial/ethnic differences.

Applications from minority-serving institutions should be judged on their capacity to conduct research rather than relying on the institutions’ research track records. In our experience, applications from institutions with strong community ties are better equipped to enroll and retain subjects in clinical and biomedical research. The importance and novelty of studies focused primarily or solely on minority populations should be recognized for their validity and worth, as these may be the only studies to recruit sufficient minority participants to determine whether research findings can be generalized to these populations.

Given the systemic bias against minority scientists, the solution does not lie in simply increasing the number of competitive applicants. To this end, the NIH is actively funding investigations to understand and eliminate discrepancies for minority investigators in the peer review process [51]. In September 2014, the NIH announced [52] winners of two competitions on increasing the fairness and impartiality of the scientific review process and for novel methods of identifying bias. A program assessing the complete anonymization of grant applications is also being piloted [53]. These efforts are part of a larger campaign to identify and root out unconscious bias in peer review [41,51]. The NIH must act on these data to ensure a just and fair voice for all stakeholders.

NIH proposals passing scientific peer review are forwarded to a second level of review, conducted by Institute and Center (IC) National Advisory Councils or Boards (henceforth referred to as “Councils”). NIH Councils make funding decisions based on the priority score and the priorities of the IC, which have varying levels of discretionary funds. A reasonable way to fund meritorious applications that reflect the diversity priorities of the ICs is to use the discretion of the Councils. Other NIH efforts to increase support for the diversity pipeline (e.g., BUILD [54]) and for diversity-related scientific initiatives are commendable, but in the absence of strong changes throughout the review process, research will continue to suffer.

## Inclusive research needs the support of the entire country

Efforts by the NIH and other agencies to address disparities in research priorities will have limited impact unless broader themes of political and economic inequality are addressed. The most important changes in our approach to science will only come when we consider inclusion and diversity important by default—not just in biomedical science, but in all aspects of society. Homogeneity in study populations will cease when racial/ethnic and socioeconomic diversity are considered socially desirable and social norms [55], be it in study populations, academic faculty, NIH study sections, or boardrooms and classrooms.

We have suggested a number of measures for the NIH to build upon the Revitalization Act. Despite the Act’s stipulation that cost not be used as justification for failure to enroll diverse populations, no discussion of new mandates for NIH-funded research can take place without addressing the crisis of declining inflation-adjusted NIH budgets. Society and patients will benefit when the NIH exercises the full scope of power provided under the 1993 Act: a call for the inclusion of historically under-represented communities in clinical research. The NIH alone will not be able to correct the disparities or inequities of the healthcare system, but it can send a powerful message that may promote changes in our health care and health science systems. There must be a collective will to prioritize diversifying our study populations, rallied by outreach to the lay community to educate voters who can exercise their franchise to their own best healthcare interests.

Fulfilling the promise of the Revitalization Act does not pit a future of precision medicine and the advancement of science against the realization of social justice for under-represented communities. Rather, the choice to study diverse populations is itself a promising path toward sound science. By reprioritizing our approach to clinical research and recruitment, we may accomplish an even greater goal: to usher in a new era of scientific discovery and health prosperity for all citizens of the world.

